# HSC71 acetylation confers protection against *Spiroplasma eriocheiris* infection by inhibiting apoptosis and promoting ROS production in arthropods

**DOI:** 10.64898/2026.03.02.708988

**Authors:** Yubo Ma, Xiang Meng, Xin Yin, Yu Yao, Siyu Lu, Wei Gu, Qingguo Meng

## Abstract

Heat shock proteins (HSP) are increasingly recognized as pivotal regulators of host innate immunity. However, the role of HSP70 post-translational modifications in modulating the immunologic functions during pathogenic infection remains poorly understood. Here, we demonstrated that heat shock cognate 71 kDa protein (HSC71), a member of the HSP70 family, was deacetylated following infection with intracellular pathogen *Spiroplasma eriocheiris*. HSC71 deficiency exacerbated hemocyte apoptosis and increased host susceptibility to *S. eriocheiris* infection. Conversely, ectopic expression of hyperacetylation-mimetic HSC71(K579Q) in *Drosophila* S2 cells conferred robust protection against *S. eriocheiris* by suppressing apoptosis and augmenting reactive oxygen species (ROS) production. Mechanistically, carnitine O-acetyltransferase (Crat) was identified as the specific acetyltransferase for HSC71. Crat acetylated HSC71 at lysine 579 (K579), which in turn impeded its interaction with the E3 ubiquitin ligase CHIP, thereby stabilizing HSC71 by attenuating ubiquitin-mediated degradation. Crat deficiency induced apoptosis and reduced ROS levels in hemocytes, thereby exacerbating *S. eriocheiris* intracellular proliferation. Furthermore, acetylation of HSC71 at K579 weakened its binding to superoxide dismutase (SOD), resulting in impaired SOD activity and consequent accumulation of microbicidal ROS. Pharmacological inhibition of the deacetylase SIRT1 with EX-527 enhanced HSC71 acetylation and ROS production, conferring resistance to *S. eriocheiris* infection in crabs. Notably, EX-527 similarly enhanced the acetylation of *Drosophila* HSC71 homolog, HSPA8, which in turn impaired its interaction with SOD. This led to elevated ROS levels and restricted intracellular proliferation of *S. eriocheiris* in S2 cells, demonstrating evolutionary conservation of this mechanism among arthropods. Therefore, this study established the modulation of HSC71 acetylation as a promising strategy to enhance arthropod immunity.

**Author summary:** Members of the HSP70 family are indispensable components of host innate immunity, and their functions are finely tuned by post-translational modifications. This study investigated the mechanisms that underlie acetylation modification of HSC71 mediated resistance to *S. eriocheiris* infection in crab. Our results revealed that the acetyltransferase Crat and the antioxidant enzyme SOD directly interacted with HSC71. Mechanistically, Crat acetylated HSC71 at K579, which prevented its ubiquitination by promoting the disassociation of E3 ubiquitin ligase CHIP, thereby improving HSC71 stability. In HSC71-deficient or Crat-deficient crabs, hemocyte apoptosis was markedly enhanced, leading to higher host mortality upon *S. eriocheiris* challenge. Meanwhile, K579 acetylation on HSC71 weakened the interaction between HSC71 and SOD, resulting in the accumulation of intracellular ROS and thereby restricting *S. eriocheiris* propagation. Administration of the SIRT1 inhibitor EX-527 enhanced HSC71 acetylation, elevated ROS production and reduced host susceptibility to *S. eriocheiris* infection. These findings revealed a novel role for Crat-mediated acetylation of HSC71 in orchestrating host defense through two synergistic mechanisms: suppressing apoptosis and promoting ROS accumulation. Thus, targeting HSC71 acetylation may represent a promising avenue to combat *S. eriocheiris* infection.

## Introduction

The heat shock protein 70 (HSP70) family comprises evolutionarily conserved yet functionally versatile molecular chaperones. They are essential for multiple proteostatic processes, including assisting in nascent protein folding, refolding misfolded proteins, disaggregating protein deposits, translocation of organellar proteins, and targeting aberrant proteins for degradation [1–3]. All members of the HSP70 family share a conserved structure comprising an N-terminal nucleotide-binding domain (NBD) and a C-terminal substrate-binding domain (SBD). The SBD consists of a β-sheet subdomain that forms the substrate-binding pocket and an α-helical lid subdomain. ATP binding and hydrolysis in the NBD drive conformational changes in the SBD, primarily through the opening and shutting of the lid, which regulates the substrate-binding affinity of HSP70 [4, 5]. Moreover, the conformational equilibrium could be precisely modulated through post-translational modifications at allosteric hotspots within its SBD, directly regulating chaperone activity while preserving structural integrity [6]. Under cellular stress conditions, upregulated HSP70 facilitates the refolding of misfold proteins to their native conformation or direct their clearance via the ubiquitin-proteasome system and chaperone-mediated autophagy, thereby mitigating cellular damage caused by aberrant proteins and alleviating associated disease pathogenesis [7]. Furthermore, HSP70 is frequently exploited by clinically important viruses (herpesvirus, retrovirus, flavivirus) to support key stages of their life cycles, particularly involving the assembly and stabilization of viral replication complexes and virion structures [8–10]. Conversely, HSP70 also contributes to host innate immunity against pathogens in aquatic animals. In teleost fish, HSP70 interacted with HMGB1b to facilitate its cytosolic translocation, which enhanced the association between HMGB1b and Beclin-1, thereby activating autophagy to restrict grass carp reovirus (GCRV) replication [11]. Similarly, shrimp HSP70 bound to Toll4, leading to activation of the transcription factor Dorsal and subsequent upregulation of antimicrobial peptides (AMPs), conferring resistance to white spot syndrome virus (WSSV) infection [12]. Therefore, elucidating the mechanism by which HSP70 regulates innate immunity is beneficial for developing novel therapies against aquatic infectious diseases.

Accumulating evidence demonstrates that post-translational modifications finely tune HSP70 function through distinct mechanisms. For instance, HSP70 methylation enhanced its binding to the Bcl2 mRNA 3’-UTR, upregulating Bcl2 protein expression, thereby promoting cancer cell survival under stress or therapeutic challenge [13]. C-terminal phosphorylation of HSP70 shifted its co-chaperone preference from CHIP to HOP, a switch that favored cellular protein folding over degradation [14]. Acetylation of HSP70 at K159 facilitated the assembly of Vps34-Beclin-1 complex and recruits E3 SUMO ligase KAP1 to promote Vps34 SUMOylation, ultimately supporting autophagosome formation [15]. Among these modifications, acetylation of lysine residues is a key post-translational modification that regulates protein function by altering conformation, subcellular localization, stability or crosstalk with other modifications, thereby influencing diverse cellular processes such as metabolism, apoptosis and innate immunity [16–19]. In breast cancer cells, HSP70 could be acetylated at K159 by p300 and deacetylated by HDAC6 [15]. Furthermore, during the early stress response in HEK293T cells, acetylation at K77 by ARD1 promoted binding to the co-chaperone HOP to facilitate refolding; subsequent deacetylation by HDAC4 switched HSP70 binding to the E3 ligase CHIP, targeting clients for degradation [20].

Notably, lysine residues are common sites for competing modifications, particularly acetylation and ubiquitination, with acetylation often antagonizing ubiquitin-proteasome-mediated degradation [21, 22]. These evidences showed that acetylation of HSP70 might regulate its stability, either by directly competing with ubiquitination at overlapping lysine residues or by modulating its interaction with specific E3 ubiquitin ligases/deubiquitinases. Despite these advances in higher vertebrates, the specific acetyltransferases or deacetylases that modified HSP70 in invertebrates, and the functional impact of such acetylation, remain largely unexplored. Reactive oxygen species (ROS) encompass a variety of derivatives of molecular oxygen that occur as a normal attribute of aerobic metabolism. These radical species can originate from exogenous resources or be generated intracellularly, primarily via the mitochondrial electron transport chain and NADPH oxidases (NOXs) [23, 24]. Elevated intracellular ROS levels exhibit diverse biological effects, ranging from oxidative damage to cellular components to the activation of specific signaling pathways [23]. In the context of host immunity, ROS function as antimicrobial effector molecules that enhance nuclear factor-κB (NF-κB) transcriptional activity and contribute to the formation of DNA-based neutrophil extracellular traps [25]. Their highly reactive nature also enables ROS to directly damage structural components of the bacterial cell membrane, thereby attenuating bacterial virulence [26]. In invertebrates, such as *Drosophila*, microbial infection induced a dramatic upregulation of dual oxidase (Duox), catalyzing the production of microbicidal ROS essential for pathogen clearance and host survival [27]. The entry of extracellular ROS into the cytosol further amplified immune responses by signaling the expression of AMPs, which acted synergistically with ROS to exert bactericidal activity [28]. Conversely, during virus infection, ROS production occurred rapidly, and the viral protein wsv220 promoted nuclear translocation of nuclear factor erythroid 2-related factor 2 (Nrf2), leading to the upregulation of antioxidant genes that mitigated ROS accumulation and facilitated WSSV replication [29]. In our previous study, infection of *Drosophila* S2 cells with intracellular pathogen *Spiroplasma eriocheiris* significantly elevated intracellular ROS levels, which peaked at 9 h post-infection and subsequently declined sharply from 12 to 72 h post-infection [30]. This dynamic modulation of ROS homeostasis prompted us to investigate its role in the pathogenesis of *S. eriocheiris*.

In this study, initial acetylome profiling showed that acetylation of HSC71 (an HSP70 family member) was downregulated following *S. eriocheiris* infection. This led us to investigate how HSC71 acetylation might influence crab antimicrobial defenses. We discovered that carnitine O-acetyltransferase (Crat) acetylated HSC71 at K579, preventing CHIP-mediated ubiquitination and degradation, thereby stabilizing HSC71 and enhancing host resistance to *S. eriocheiris* infection. In parallel, treatment with the SIRT1 inhibitor EX-527 enhanced HSC71 acetylation, which promoted its dissociation from superoxide dismutase (SOD) and elevated intracellular ROS levels, leading to protection against *S. eriocheiris* infection in both crabs and *Drosophila* S2 cells. Collectively, our findings identified Crat-mediated acetylation of HSC71 as a novel regulatory mechanism that orchestrated innate immune responses.

## Results

### Protective effect of HSC71 against *S. eriocheiris* infection in crab

Acetylation is a significant post-translational modification involved in host antipathogen responses [16]. To examine acetylated proteins critical for defense against *S. eriocheiris*, we performed acetylome analysis of hemocytes from healthy and infected crabs. Compared with control groups, we identified 79 peptides with upregulated acetylation and 107 peptides with downregulated acetylation following infection, using a cutoff of |fold change| > 2 and adjusted *p*-value < 0.05 (Fig 1A, 1B; S1 Fig). Notably, acetylation of HSC71 was significantly reduced upon infection (0.137-fold change, *p*-value=0.011) (S1 Table). Consistent with these findings, IP with an anti-pan-acetyllysine antibody confirmed that *S. eriocheiris* infection markedly reduced HSC71 acetylation in hemocytes (Fig 1C). Sequence analysis revealed that HSC71 possessed an open reading frame of 1953 bp, encoding a 651-amino-acid protein with a predicted molecular weight of 71.27 kDa, and exhibited high sequence similarity throughout vertebrates and invertebrates (S2 Fig). Upon *S. eriocheiris* infection, the mRNA level of HSC71 was significantly downregulated early after infection but upregulated at later stages (Fig 1D). To assess the function of HSC71 during *S. eriocheiris* infection, HSC71 was silenced by specific siRNA in crabs which were subsequently challenged with *S. eriocheiris*. Quantitative real-time PCR and western blot revealed significant reductions in both mRNA and protein levels of HSC71 in hemocytes (Fig 1E). Meanwhile, HSC71 deficiency significantly promoted hemocyte apoptosis, indicated by increased Annexin V-positive cells and loss of mitochondrial membrane potential (Fig 1F, 1G). Importantly, HSC71-deficient crabs exhibited increased susceptibility to *S. eriocheiris* infection, as evidenced by elevated *S. eriocheiris* load in hemocytes and reduced survival compared to controls (Fig 1H, 1I). These results indicated that HSC71 was required for crab resistance to *S. eriocheiris* infection.

**Fig. 1.**
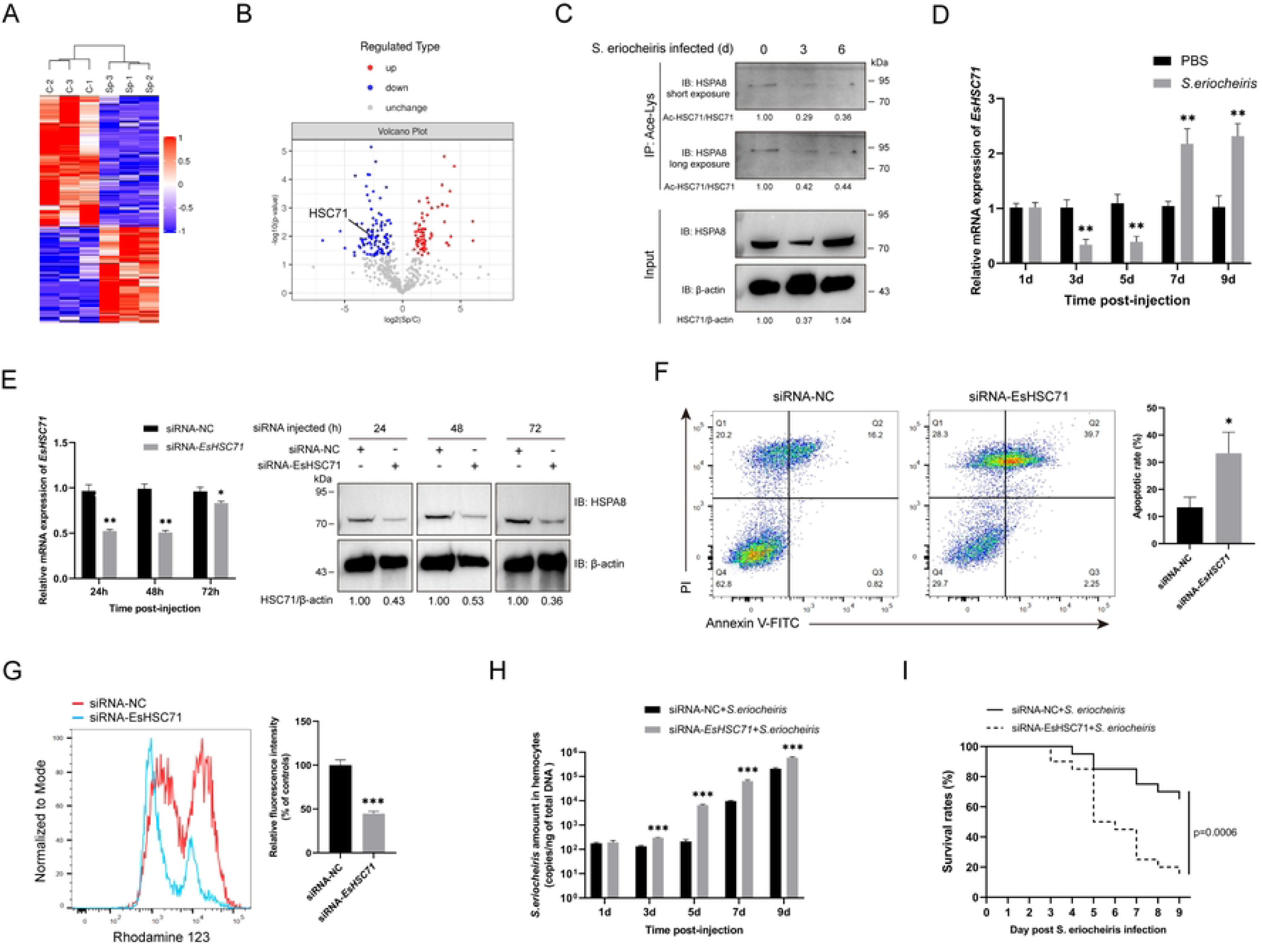
HSC71 protects crabs from *S. eriocheiris* infection. (A) Heatmap showing the differentially acetylated proteins in *S. eriocheiris* infection. (B) Volcano plot of the differentially acetylated proteins in *S. eriocheiris* infection. (C) Acetylation of endogenous HSC71 in hemocytes at various times post *S. eriocheiris* infection. Acetylation levels of HSC71 were assessed by IP with an anti-Acetyllysine antibody, followed by immunoblotting with an anti-HSPA8 antibody. (D) Transcription profiles of HSC71 in hemocytes were detected by quantitative real-time PCR at various intervals post-infection with *S. eriocheiris*. (E) Knockdown of HSC71 was validated by quantitative real-time PCR (left) and western blot (right). (F) Flow cytometric analysis of apoptosis in crabs hemocytes following 24 h siRNA treatment and Annexin V-FITC/PI labeling. The graph on the right showed the apoptosis rate, defined by the percentage of hemocytes in the Q2 and Q3 quadrants. (G) Flow cytometric analysis of mitochondrial membrane potential with Rhodamine 123 in hemocytes from siRNA-treated crabs. The graph on the right showed the quantification of the mean fluorescence intensity ratio of Rhodamine 123. (H) Absolute quantitative PCR analysis of the *S. eriocheiris* copy number in hemocytes of HSC71-deficient crabs in *S. eriocheiris* infection. (I) Survival analysis of HSC71-deficient crabs following *S. eriocheiris* infection. Crabs were pretreated with siRNA for 24 h before *S. eriocheiris* infection and monitored daily for survival. Statistical significance was determined by the Log rank test (*n*=20). Statistical analysis (D, E, F, G and H) was performed using the two-tailed student’s t-test (\**P*<0.05, \*\**P*<0.01, \*\*\**P*<0.001). Data represented the mean ± SD of triplicate assays.

### Lysine 579 acetylation of HSC71 protects *Drosophila* S2 cells from *S. eriocheiris* infection by inhibiting apoptosis

To investigate the role of HSC71 acetylation in *S. eriocheiris* pathogenesis, we first identified a potential acetylation site at lysine 579 (K579) based on our previous acetylome data (Fig 2A; S1 Table). Multiple sequence alignment showed that K579 was less conserved in *Homo sapiens*, *Drosophila melanogaster* or *Danio rerio*, but was strictly conserved in crustaceans, including *Penaeus vannamei*, *Penaeus japonicus*, and *Scylla paramamosain* (Fig 2B). The lysine 579 residue was mutated to glutamine (K579Q) to mimic the hyperacetylated state or to arginine (K579R) to mimic the hypoacetylated state (Fig 2C). These mutant constructs were then overexpressed successfully in *Drosophila* S2 cells for functional analysis (Fig 2D). Following *S. eriocheiris* infection, the acetylation level of ectopically expressed HSC71 was significantly reduced, and this reduction was blocked by the K579Q mutation, confirming K579 as a major site for acetylation during infection (Fig 2E). Flow cytometry revealed that K579Q mutation significantly inhibited cellular apoptosis upon *S. eriocheiris* infection (Fig 2F). Consistent with this, cells expressing K579Q mutant exhibited markedly increased viability and a drastically reduced *S. eriocheiris* load (Fig 2G, 2H). Altogether, HSC71 acetylation at K579 was confirmed to beneficial for host defense against *S. eriocheiris* infection.

**Fig. 2.**
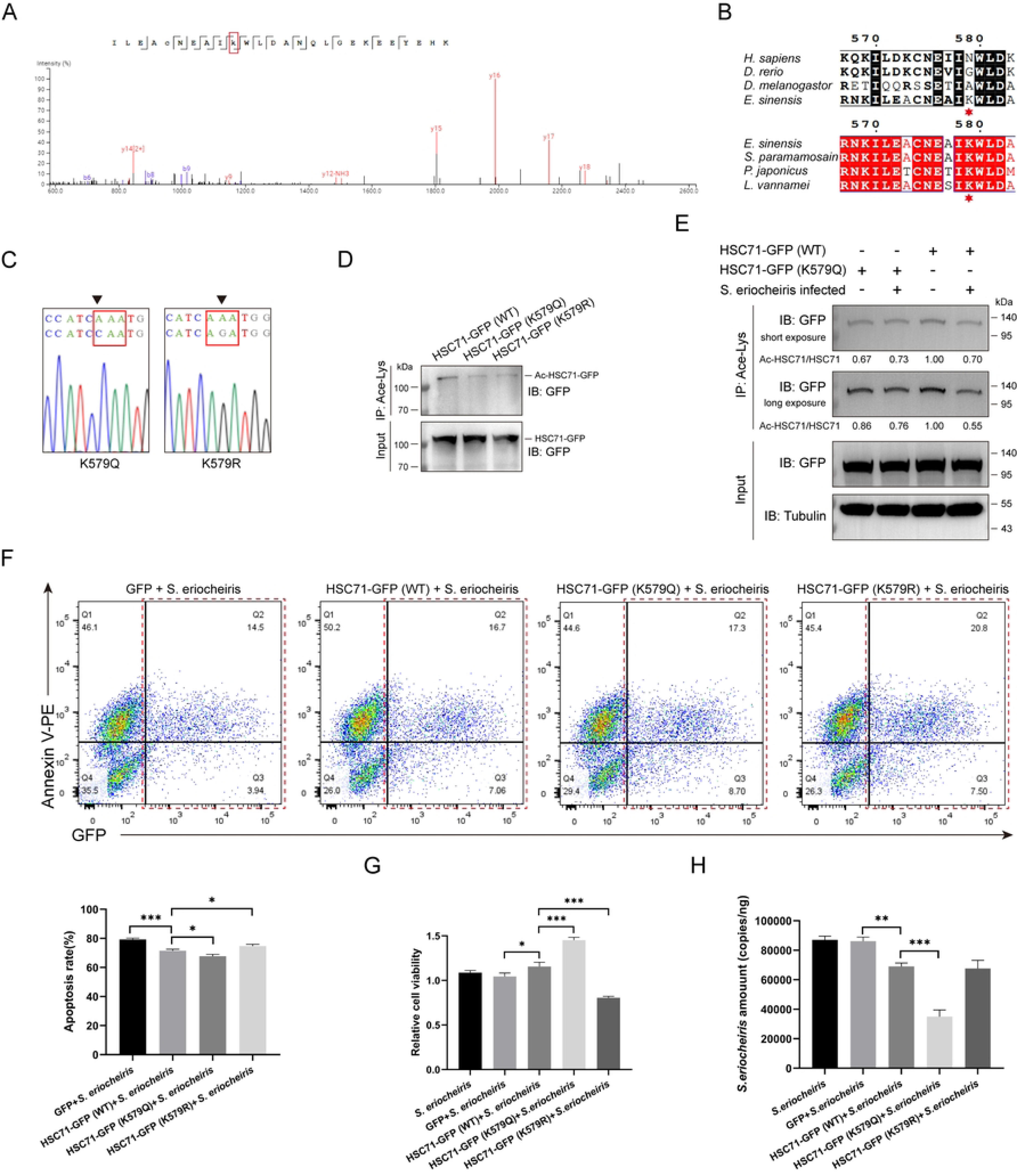
Lysine 579 acetylation of HSC71 attenuates *S. eriocheiris*-induced apoptosis and reduces the proliferation of *S. eriocheiris*. (A) Identification of lysine acetylation at residue 579 (K579) by mass spectrometry. (B) Sequence alignment of the K579 acetylation site in crab HSC71 with homologs from other species. The K579 residues were marked by red asterisks. (C) DNA sequencing results of mutant plasmids expressing HSC71-GFP (K579Q) and HSC71-GFP (K579R). (D) Acetylation of ectopically expressed HSC71 in *Drosophila* S2 cells. Lysates from cells overexpressing wild-type HSC71-GFP or mutants were immunoprecipitated with an anti-Acetyllysine antibody, followed by immunoblotting with an anti-GFP antibody to assess the acetylation status of exogenous HSC71. (E) *S. eriocheiris* infection induced deacetylation of HSC71 at K579 in *Drosophila* S2 cells. Cells overexpressing wild-type HSC71-GFP or mutant (K579Q) were infected with *S. eriocheiris* for 24 h or uninfected. The acetylation status of exogenous HSC71 was analyzed by IP and immunoblotting as described above. (F) Apoptosis analysis of *Drosophila* S2 cells transfected with the indicated plasmids and subsequently infected with *S. eriocheiris* for 48 h was detected by flow cytometry with Annexin V-PE labeling. Apoptotic rates were analyzed in gated GFP-positive (transfected) cells from Q2 and Q3 quadrants, calculated as the percentage of cells in Q2 relative to the total in Q2 and Q3 (bottom). (G) Cell viability of *Drosophila* S2 cells transfected with the indicated plasmids was evaluated by the CCK-8 assay after *S. eriocheiris* infection for 48 h. (H) The copy number of *S. eriocheiris* in *Drosophila* S2 cells transfected with the indicated plasmids was quantified using absolute quantitative PCR after *S. eriocheiris* infection for 48 h. For (F, G and H), the two-tailed unpaired student’s t-test was used for statistical analysis (\**P*<0.05, \*\**P*<0.01, \*\*\**P*<0.001). Data represented the mean ± SD of triplicate assays.

### Identification of HSC71-interacting proteins

To investigate the mechanism by which HSC71 acetylation conferred protection against *S. eriocheiris* infection, IP was performed on hemocytes using anti-HSPA8 antibody to isolate HSC71-interacting proteins, with mouse IgG as a control for non-specific binding. Coomassie staining revealed distinct differential bands, and subsequent mass spectrometry analysis identified 25 candidate proteins that specifically Co-immunoprecipitated with HSC71 (Fig 3A; S2 Table). Among the candidate interacting proteins, Crat is of our special interest due to its potential, as an acetyltransferase, to regulate HSC71 acetylation. Based on this, we confirmed the interaction between HSC71 and Crat by Co-immunoprecipitation (Co-IP) following co-transfection of HSC71-HA and Crat-GFP plasmids into S2 cells (Fig 3B). Furthermore, overexpression of Crat-GFP significantly enhanced the acetylation of exogenous HSC71-HA (Fig 3C), indicating that Crat is a specific acetyltransferase for HSC71. Moreover, mass spectrometry identified several antioxidant proteins as putative HSC71 interactors. We focused on SOD and peroxiredoxin due to their roles in eliminating ROS, which is crucial for host defense against pathogens. Co-IP assays confirmed that HSC71-HA specifically interacted with either SOD-GFP or Prx-GFP when co-expressed in *Drosophila* S2 cells (Fig 3D, 3E). Laser confocal microscopy further revealed the colocalization of HSC71-GFP with Crat-RFP, SOD-RFP and Prx-RFP in the cytosol (Fig 3F), suggesting that HSC71 acetylation might modulate ROS homeostasis as part of the host defense against *S. eriocheiris* infection.

**Fig. 3.**
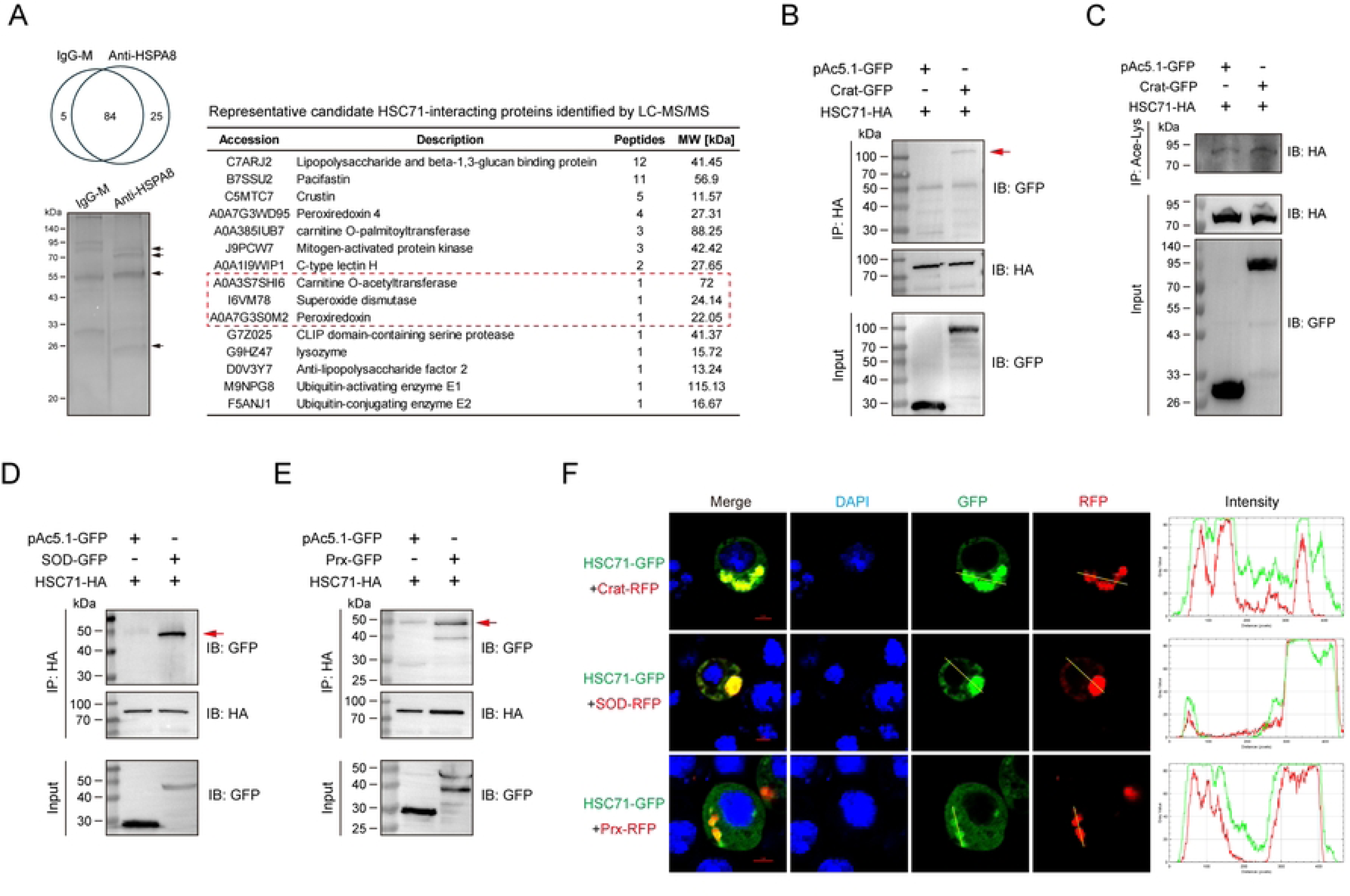
HSC71 interacts with Crat, SOD and Prx. (A) IP analysis of HSC71-interacting proteins from hemocytes using an anti-HSPA8 antibody, with mouse IgG serving as a control to exclude non-specific binding proteins. The precipitates were analyzed by Coomassie staining and mass spectrometry. Differential bands were indicated by black arrows. The graph on the right summarized representative candidate HSC71-interacting proteins identified by mass spectrometry. (B) Interaction between Crat and HSC71 was validated by Co-IP. *Drosophila* S2 cells were co-transfected with HSC71-HA, together with Crat-GFP or GFP plasmids for 48 h. Cell lysates were immunoprecipitated with an anti-HA antibody, followed by immunoblotting with an anti-GFP antibody. (C) Effect of Crat on HSC71 acetylation. Cells were transfected as indicated above. The acetylation level of HSC71-HA in *Drosophila* S2 cells was assessed by IP with an anti-Acetyllysine antibody, followed by immunoblotting with an anti-HA antibody. (D and E) Validation of interactions between HSC71 and SOD (D) or Prx (E) by Co-IP. Lysates from *Drosophila* S2 cells co-expressing the indicated plasmids were immunoprecipitated with an anti-HA antibody, and the precipitates were analyzed by immunoblotting with an anti-GFP antibody. For (B, D and E), red arrows indicated bands corresponding to HSC71-interacting proteins. (F) Colocalization of HSC71 with Crat, SOD and Prx in *Drosophila* S2 cells. Cells were co-transfected with HSC71-GFP, together with Crat-RFP, SOD-RFP or Prx-RFP plasmids for 48 h. Protein colocalization was visualized by laser confocal microscopy. Scale bar, 2 μm.

### Acetylation of HSC71 at K579 by Crat controls HSC71 stabilization to limit *S. eriocheiris* infection

To identify the acetyltransferase responsible for HSC71 acetylation, we assessed the acetylation level of endogenous HSC71 following Crat knockdown in crabs. Quantitative real-time PCR showed that *S. eriocheiris* infection led to downregulation of Crat in hemocytes, which aligned with a concurrent reduction in HSC71 acetylation (Fig 4A). Employing siRNA specifically targeted for Crat, the mRNA level of Crat was significantly reduced in hemocytes (Fig 4B). Subsequently, we observed a notable reduction in HSC71 acetylation and protein levels in hemocytes following siRNA-Crat application (Fig 4C). However, Crat knockdown did not alter the mRNA level of HSC71 (Fig 4D), indicating that HSC71 stability might be regulated at the post-translational level. Through co-expression of Crat and wild-type and mutant of HSC71, we found that Crat significantly promoted the acetylation and stabilization of HSC71 in *Drosophila* S2 cells, whereas the increase in acetylation and protein levels was markedly abolished by the mutation K579 to arginine (Fig 4E). Moreover, Crat overexpression reduced the ubiquitination level of HSC71 (Fig 4F), suggesting that Crat-mediated acetylation of HSC71 at K579 might counteract its ubiquitination to regulate protein stability. Given the well-recognized role of CHIP in ubiquitin-proteasomal degradation of HSC70 proteins in vertebrates [31], we tested whether Crat stabilized HSC71 by interfering with this process. In *Drosophila* S2 cells, we found that CHIP could strongly bind to HSC71. However, this interaction was evidently reduced in the presence of Crat (Fig 4G). Furthermore, CHIP-mediated HSC71 ubiquitination was largely abrogated by Crat overexpression (Fig 4H). We next examined the ubiquitination levels of wild-type and mutant of HSC71. The hyperacetylation-mimetic HSC71(K579Q) exhibited reduced ubiquitination in *Drosophila* S2 cells (Fig 4I). In agreement with this, the interaction of CHIP and HSC71 was significantly attenuated by the mutation K579 to glutamine (Fig 4J). Meanwhile, co-expression of CHIP reduced the protein level of ectopically expressed HSC71 in *Drosophila* S2 cells (Fig 4K). However, Crat overexpression could reverse the reduction in HSC71 protein level induced by CHIP (Fig 4L). These results indicated that Crat competitively bound to HSC71, thereby antagonizing CHIP-mediated HSC71 ubiquitination and degradation. To dissect the role of Crat in immune regulation, crabs were administered with Crat-specific siRNA prior to *S. eriocheiris* infection. As expected, Crat knockdown significantly increased HSC71 ubiquitination and protein levels in hemocytes (Fig 4M). Flow cytometry revealed that Crat deficiency exacerbated hemocyte apoptosis and loss of mitochondrial membrane potential (Fig 4N, 4O). Further, Crat knockdown significantly elevated *S. eriocheiris* load in hemocytes, and reduced crab survival following infection (Fig 4P, 4Q). These results demonstrated that Crat contributed to host defense against *S. eriocheiris* infection by acetylating and stabilizing HSC71.

**Fig. 4.**
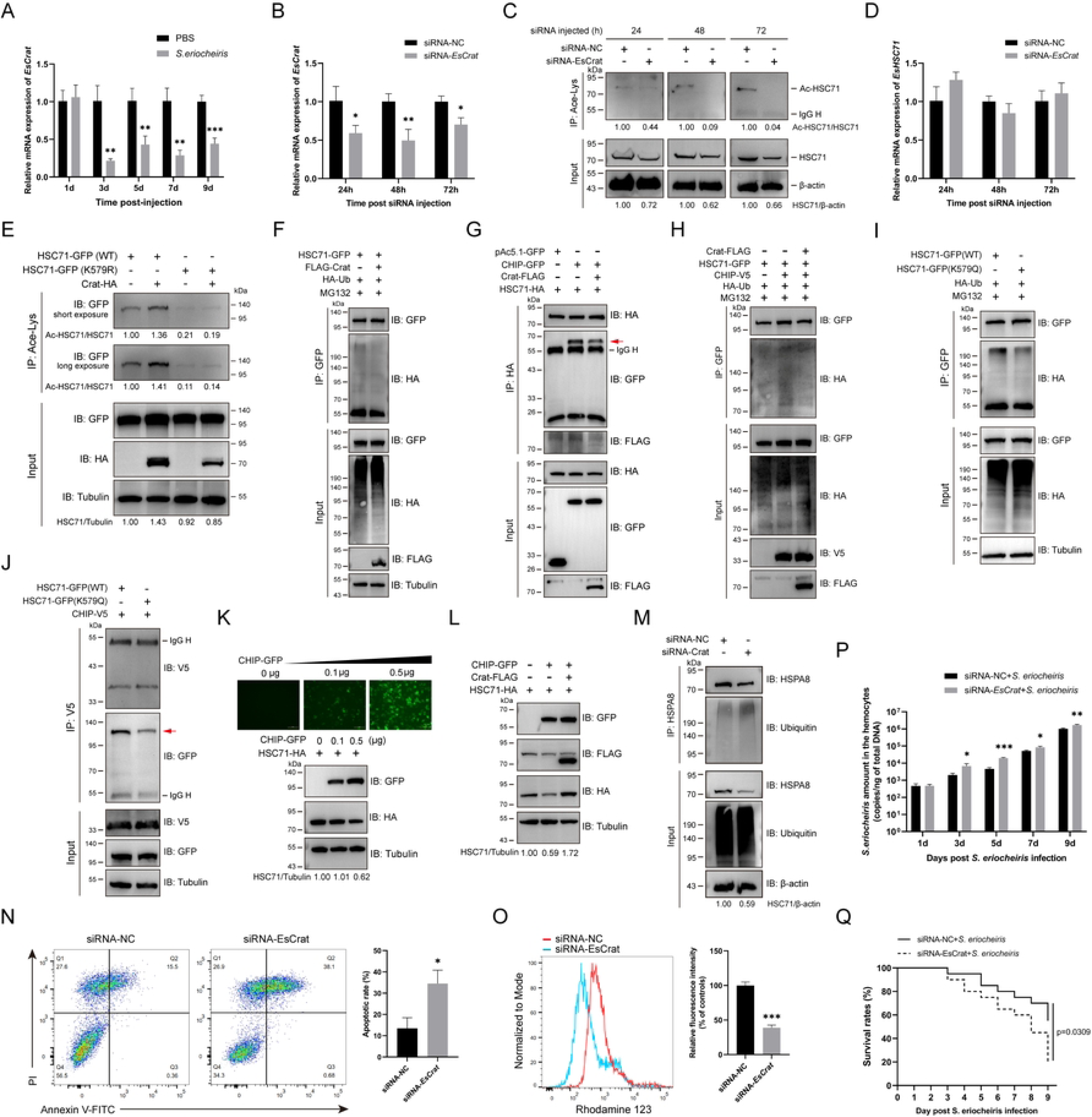
Crat acetylates HSC71 at K579 and stabilizes HSC71 via impairing CHIP-mediated ubiquitination of HSC71 to restrict *S. eriocheiris* replication. (A) Transcription profiles of Crat in hemocytes were detected by quantitative real-time PCR post-infection with *S. eriocheiris*. (B) Knockdown of Crat was validated by quantitative real-time PCR. (C) Effect of Crat knockdown on HSC71 acetylation. At 24, 48 and 72 h post siRNA injection, lysates from hemocytes were immunoprecipitated with an anti-Acetyllysine antibody, followed by immunoblotting with an anti-HSC71 antibody. (D) HSC71 transcription levels after Crat knockdown was determined by quantitative real-time PCR. (E) Crat acetylated HSC71 at K579. *Drosophila* S2 cells were co-transfected with plasmids expressing wild-type HSC71-GFP or mutant (K579R) and empty vector (HA) or Crat-HA. The acetylation level of HSC71 was assessed by IP with an anti-Acetyllysine antibody, followed by immunoblotting with an anti-GFP antibody. (F) The ubiquitination level of HSC71 after Crat overexpression. *Drosophila* S2 cells were co-transfected with plasmids expressing HSC71-GFP, HA-Ub and empty vector (FLAG) or FLAG-Crat. After treatment with MG132 (20 μM) for 8 h, HSC71-GFP was immunoprecipitated with anti-GFP magnetic beads, and the precipitates were analyzed by immunoblotting with an anti-HA antibody. (G) Effect of Crat on the interaction between CHIP and HSC71. *Drosophila* S2 cells were co-transfected with plasmids expressing HSC71-HA and empty vector (GFP) or CHIP-GFP, together with empty vector (FLAG) or Crat-FLAG. IP was performed using an anti-HA antibody, followed by immunoblotting with anti-GFP and anti-FLAG antibodies. (H) Effect of Crat on CHIP-mediated ubiquitination of HSC71. *Drosophila* S2 cells co-expressing the indicated plasmids were treated with MG132 (20 μM) for 8 h before harvesting. IP was performed using anti-GFP magnetic beads, followed by immunoblotting with an anti-HA antibody. (I) Effect of K579 acetylation on HSC71 ubiquitination. *Drosophila* S2 cells co-expressing wild-type HSC71-GFP or mutant (K579Q) and HA-Ub were treated with MG132 (20 μM) for 8 h before harvesting. Ubiquitination levels were assessed by IP with anti-GFP magnetic beads and immunoblotting with an anti-HA antibody. (J) Effect of K579 acetylation on the interaction between HSC71 and CHIP. *Drosophila* S2 cells were co-transfected with plasmids expressing wild-type HSC71-GFP or mutant (K579Q) and CHIP-V5. Co-IP was performed using an anti-V5 antibody, followed by immunoblotting with an anti-GFP antibody. (K) Protein levels of HSC71 after CHIP overexpression in *Drosophila* S2 cells. Cells were co-transfected with HSC71-HA, together with a dose gradient of CHIP-GFP plasmids for 72 h. Overexpression of CHIP was validated by fluorescence microscopy and immunoblotting with an anti-GFP antibody. HSC71 protein levels were analyzed by immunoblotting with anti-HA and anti-Tubulin antibodies. (L) Effect of Crat overexpression on HSC71 protein levels in the presence of CHIP. *Drosophila* S2 cells were co-transfected with the indicated plasmids for 72 h. HSC71 protein levels were analyzed by immunoblotting with anti-HA and anti-Tubulin antibodies. (M) Effect of Crat deficiency on HSC71 ubiquitination. Crab hemocytes were isolated 24 h after siRNA injection. Ubiquitination levels were assessed by IP with an anti-HSPA8 antibody and immunoblotting with an anti-Ubiquitin antibody. (N) Effect of Crat deficiency on hemocyte apoptosis. Crabs were treated with siRNA for 24 h, and hemocyte apoptosis was analyzed by flow cytometry using Annexin V-FITC/PI staining. The right histogram represented the percentage of apoptotic cells in Q2 and Q3 quadrants. (O) Effect of Crat deficiency on mitochondrial membrane potential in hemocytes. Following 24 h of siRNA treatment, mitochondrial membrane potential was assessed by flow cytometry using Rhodamine 123 staining. The right histogram represented the mean fluorescence intensity ratio of Rhodamine 123. (P) Quantification of *S. eriocheiris* load in hemocytes of Crat-deficient crabs following infection with *S. eriocheiris*. Crabs were pretreated with siRNA for 24 h before challenge with *S. eriocheiris*. At the indicated times post-infection, the copy number of *S. eriocheiris* was quantified by absolute quantitative PCR. (Q) Survival analysis of Crat-deficient crabs following infection with *S. eriocheiris*. Statistical significance was determined by the Log rank test (n=20). For (A, B, D, N, O and P), the two-tailed unpaired student’s t-test was used for statistical analysis (\**P*<0.05, \*\**P*<0.01, \*\*\**P*<0.001). Data represented the mean ± SD of triplicate assays.

### Acetylation of HSC71 at K579 induces ROS-mediated antimicrobial defense in response to *S. eriocheiris* infection

To validate the hypothesis that HSC71 acetylation modulated ROS homeostasis during *S. eriocheiris* infection, the intracellular ROS levels in *Drosophila* S2 cells overexpressing wild-type HSC71-GFP or mutants were examined. Following *S. eriocheiris* infection, cells overexpressing the hyperacetylation-mimetic mutant (K579Q) exhibited a significant increase in ROS production (Fig 5A). To elucidate the mechanism underlying HSC71 acetylation-primed increase in ROS, the interaction between these HSC71 variants and antioxidant proteins were examined via Co-IP. Hyperacetylation of K579 on HSC71 (K579Q) significantly attenuated the binding of HSC71 to SOD, whereas hypoacetylation of K579 on HSC71 (K579R) showed a robust interaction (Fig 5B). The association of HSC71 with peroxiredoxin did not differ significantly between the K579Q and K579R mutants (Fig 5C), suggesting their interaction might be acetylation-independent. Similarly, the interaction between HSC71(K579Q) and DmSOD in *Drosophila* S2 cells was strikingly decreased, while the K579R mutation enhanced the interaction (Fig 5D). Moreover, SOD activity was significantly reduced in the K579Q group but increased in the K579R group (Fig 5E), suggesting acetylation at K579 impairs antioxidant capacity by disrupting the HSC71-SOD interaction, thereby promoting ROS accumulation.

**Fig. 5.**
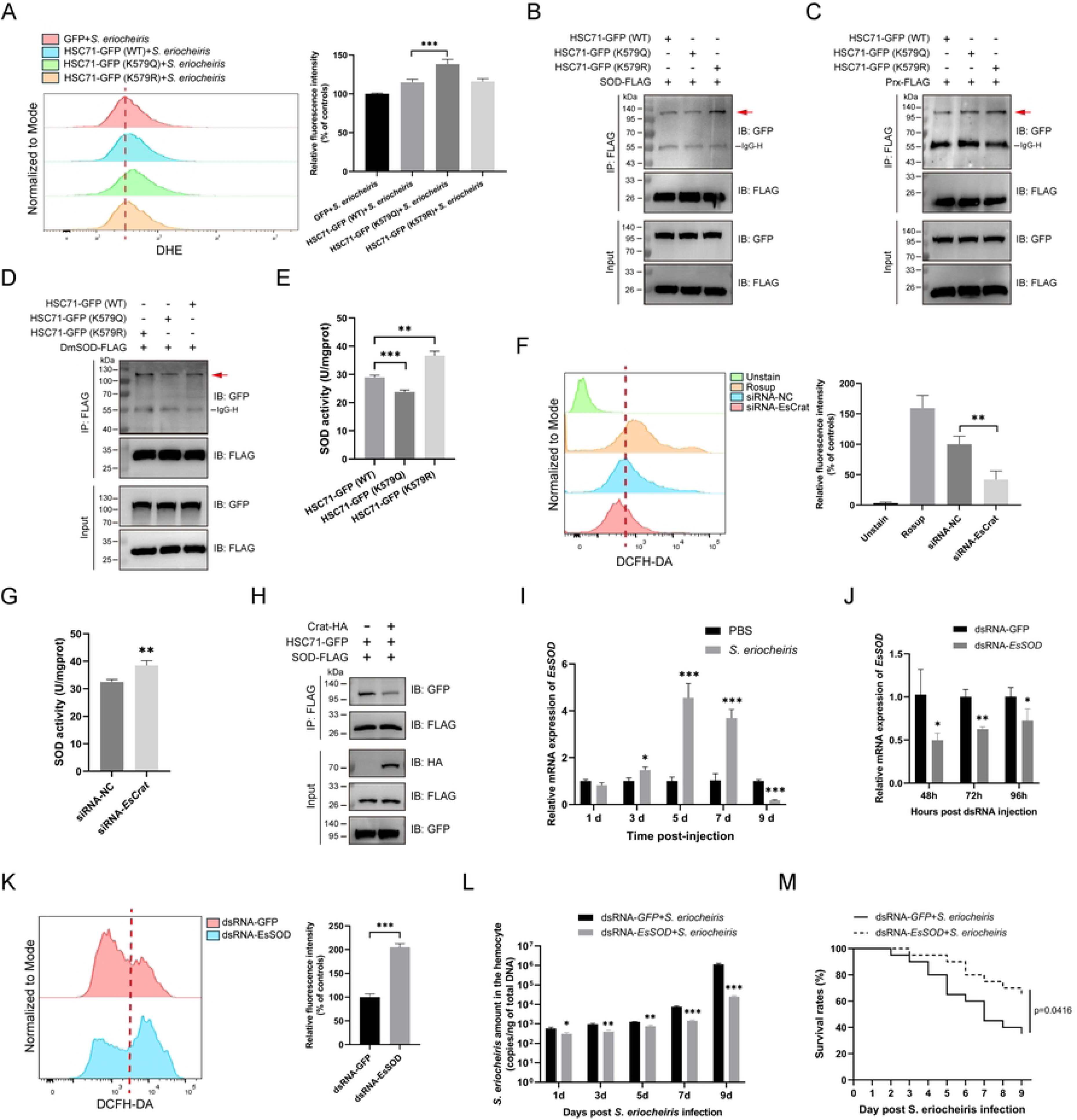
HSC71 acetylation impairs SOD binding and elevates intracellular ROS levels to augment host defense against *S. eriocheiris* (A) Intracellular ROS levels in *Drosophila* S2 cells were measured by flow cytometry. Cells overexpressing the indicated plasmids were infected with *S. eriocheiris* for 9 h and stained with DHE. The right histogram showed the quantification of the mean fluorescence intensity of DHE. (B-D) Effect of K579 acetylation on the interaction between HSC71 and SOD (B), Prx (C) and DmSOD (D). *Drosophila* S2 cells were co-transfected with plasmids expressing wild-type HSC71-GFP or mutants and empty vector (FLAG), SOD-FLAG (B), Prx-FLAG (C) or DmSOD-FLAG (D). Co-IP was performed using anti-FLAG magnetic beads, and the precipitates were analyzed by immunoblotting with an anti-GFP antibody. (E) SOD activities of *Drosophila* S2 cells overexpressing wild-type HSC71-GFP or mutants. (F) Flow cytometric analysis of intracellular ROS levels in hemocytes after Crat knockdown using DCFH-DA staining. Hemocytes from healthy crabs treated with 50 μg/mL Rosup for 30 min served as positive control. The mean fluorescence intensity of DCFH-DA was shown in the right histogram. (G) SOD activities of hemocytes after Crat knockdown. (H) Effect of Crat overexpression on the interaction between HSC71 and SOD. *Drosophila* S2 cells co-expressing the indicated plasmids were immunoprecipitated with anti-FLAG magnetic beads, and the precipitates were analyzed by immunoblotting with an anti-GFP antibody. (I) Transcription profiles of SOD in hemocytes were detected by quantitative real-time PCR post-infection with *S. eriocheiris*. (J) Knockdown of HSC71 was validated by quantitative real-time PCR. (K) Flow cytometric analysis of intracellular ROS levels in hemocytes after SOD knockdown using DCFH-DA staining. The mean fluorescence intensity of DCFH-DA was shown in the right histogram. (L) *S. eriocheiris* load in hemocytes of SOD-deficient crabs. The copy number of *S. eriocheiris* was quantified by absolute quantitative PCR at the indicated times. (M) Survival analysis of SOD-deficient crabs following infection with *S. eriocheiris*. Crabs were pretreated with dsRNA for 48 h before challenge with *S. eriocheiris* and monitored daily for survival. Statistical significance was determined by the Log rank test (n=20). Statistical analysis (A, E, F, G, I, J, K and L) was performed using the two-tailed student’s t-test (\**P*<0.05, \*\**P*<0.01, \*\*\**P*<0.001). Data represented the mean ± SD of triplicate assays.

To confirm whether HSC71 acetylation was relevant to ROS production in crabs, acetylation of endogenous HSC71 was inhibited by siRNA-Crat application. In Crat-deficient crabs, hemocytes exhibited a significant reduction in intracellular ROS levels and a concurrent increase in SOD activity (Fig 5F, 5G). IP analysis revealed that the association of HSC71 with SOD was markedly disrupted by Crat overexpression (Fig 5H). We next examined the role of SOD in ROS homeostasis and found that *S. eriocheiris* infection significantly induced SOD transcription in hemocytes (Fig 5I). Knockdown of SOD with specific dsRNA significantly reduced its transcript levels and increased intracellular ROS levels in hemocytes (Fig 5J, 5K). Furthermore, SOD silencing significantly reduced the *S. eriocheiris* load in hemocytes and improved the survival rate of infected crabs (Fig 5L, 5M), demonstrating the critical role of the ROS system in host defense against *S. eriocheiris* infection.

### Administration of EX-527 enhances HSC71 acetylation and ROS generation to resist *S. eriocheiris* infection

To further explore the acetylation of HSC71, crab hemocytes were treated with TSA (a broad-spectrum HDAC inhibitor) and NAM (a broad-spectrum SIRT inhibitor), respectively. The immunoprecipitated acetylated HSC71 was significantly increased under NAM treatment, indicating that HSC71 deacetylation was controlled by SIRT family deacetylases (Fig 6A). Using selective inhibitors, we found that SIRT1 inhibition (EX-527), but not SIRT2 inhibition (SirReal2), significantly enhanced endogenous HSC71 acetylation in crab hemocytes within 6 h, implicating SIRT1 as the primary deacetylase for HSC71 (Fig 6B). Additionally, administration of EX-527 notably elevated the intracellular ROS level and inhibited the SOD activity in hemocytes (Fig 6C, 6D). To further assess the immunoprotective effect of EX-527, crabs were administrated with EX-527 ahead of *S. eriocheiris* infection. Absolute quantitative PCR analysis revealed a significant lower *S. eriocheiris* load in hemocytes of crabs treated with EX-527 compared to the DMSO-treated control group (Fig 6E). Importantly, EX-527 significantly prolonged the survival of infected crabs (Fig 6F), demonstrating its efficacy in alleviating the pathogenesis of *S. eriocheiris* infection.

**Fig. 6.**
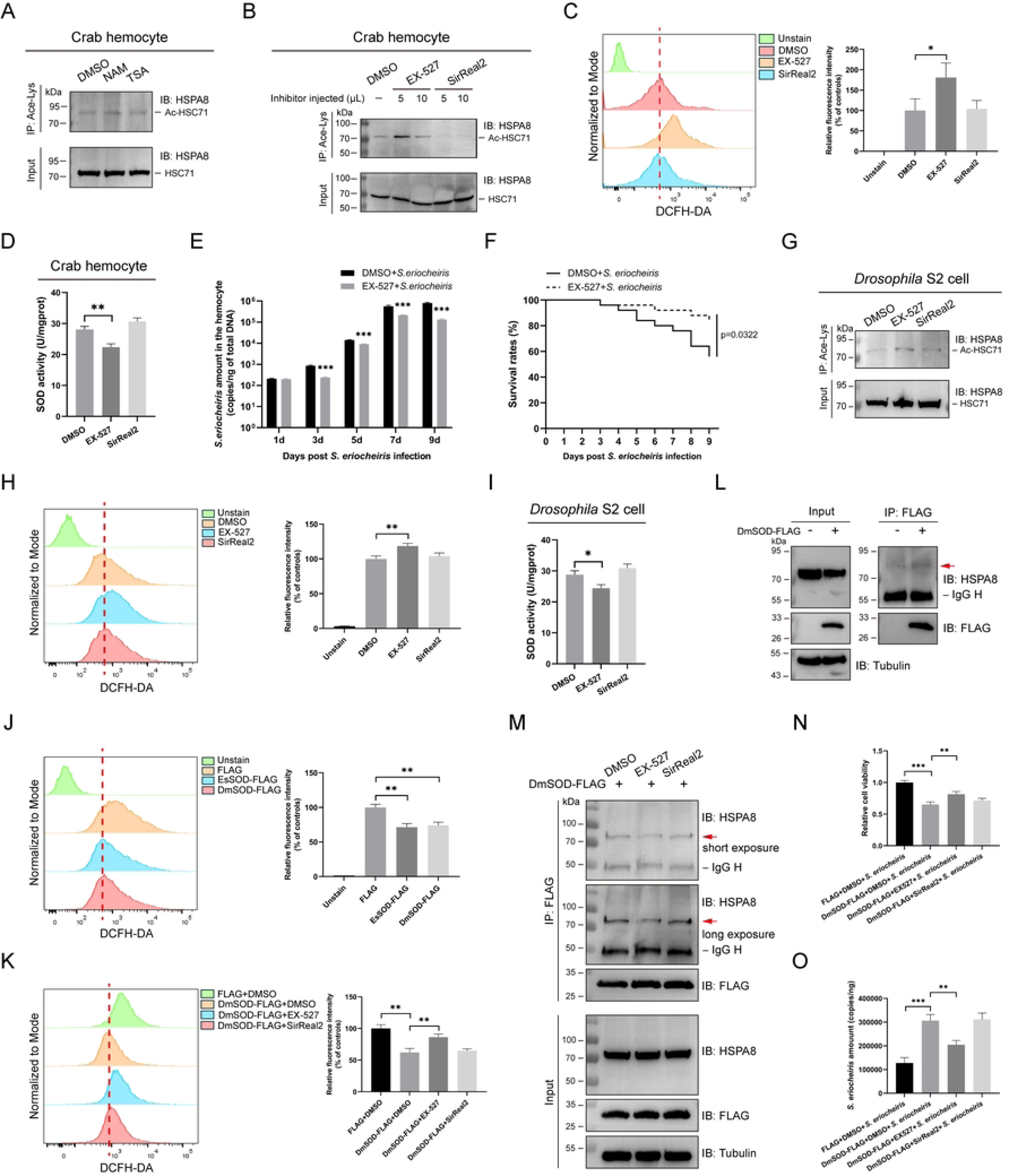
EX-527 potentiates HSC71 acetylation and ROS production to restrict *S. eriocheiris* infection. (A) Acetylation of endogenous HSC71 in hemocytes treated with deacetylase inhibitors. Primary hemocytes were treated with NAM (5 mM) or TSA (3 μM) for 12 h. HSC71 acetylation was analyzed by IP with an anti-Acetyllysine antibody, followed by immunoblotting with an anti-HSPA8 antibody. (B) Effect of SIRT inhibitors on endogenous HSC71 acetylation in hemocytes. Hemocytes from crabs injected with EX-527 or SirReal2 for 6 h were immunoprecipitated with an anti-Acetyllysine antibody, and the precipitates were analyzed by immunoblotting with an anti-HSPA8 antibody. (C) Effect of SIRT inhibitors on intracellular ROS levels in hemocytes. Hemocytes isolated from the treated crabs were analyzed by flow cytometry after staining with DCFH-DA. Quantitative analysis of the mean fluorescence intensity was shown in the right histogram. (D) Effect of SIRT inhibitors on SOD activities of hemocytes. (E) *S. eriocheiris* load in hemocytes after EX-527 treatment. The copy number of *S. eriocheiris* was quantified by absolute quantitative PCR at the indicated times. (F) Survival analysis of crabs treated with EX-527 and infected with *S. eriocheiris*. Statistical significance was determined by the Log rank test (n=25). (G) Acetylation of endogenous HSPA8 in *Drosophila* S2 cells treated with SIRT inhibitors. Cells were treated with EX-527 (1 μM) or SirReal2 (1 μM) for 24 h before harvesting. IP was performed using an anti-Acetyllysine antibody, followed by immunoblotting with an anti-HSPA8 antibody. (H) Effect of SIRT inhibitors on intracellular ROS levels in *Drosophila* S2 cells. ROS levels were assessed by flow cytometry using DCFH-DA staining, with quantitative analysis shown in the right histogram. (I) Effect of SIRT inhibitors on SOD activities of *Drosophila* S2 cells. (J) Flow cytometric analysis of intracellular ROS levels in *Drosophila* S2 cells overexpressing EsSOD or DmSOD. At 48 h post-transfection, cells were stained with DCFH-DA and the mean fluorescence intensity was quantified in the right histogram. (K) Effect of SIRT inhibitors on intracellular ROS levels in *Drosophila* S2 cells overexpressing DmSOD. Following transfection and inhibitor treatment, cells were stained with DCFH-DA for flow cytometry analysis. The quantification of the mean fluorescence intensity was shown in the right histogram. (L) Interaction between endogenous HSPA8 and DmSOD was validated by IP. (M) Effect of SIRT inhibitors on the interaction between endogenous HSPA8 and DmSOD. *Drosophila* S2 cells overexpressing DmSOD-FLAG were treated with EX-527 (1 μM) or SirReal2 (1 μM) for 24 h. The protein interaction was analyzed by IP using anti-FLAG magnetic beads, followed by immunoblotting with an anti-HSPA8 antibody. (N) Effect of SIRT inhibitors on cell viability of *Drosophila* S2 cells after *S. eriocheiris* infection. Cells transfected with DmSOD-FLAG plasmids were pretreated with EX-527 or SirReal2 before challenge with *S. eriocheiris* for 48 h. Cell viability was evaluated by the CCK-8 assay at 48 h post-infection. (O) Effect of SIRT inhibitors on *S. eriocheiris* proliferation in *Drosophila* S2 cells. Following transfection and inhibitor treatment, *Drosophila* S2 cells were infected with *S. eriocheiris* for 48 h. The copy number of *S. eriocheiris* was quantified by absolute quantitative PCR. Statistical analysis (C, E, H, I, K, L, N and O) was performed using the two-tailed student’s t-test (\**P*<0.05, \*\**P*<0.01, \*\*\**P*<0.001). Data represented the mean ± SD of triplicate assays.

To determine whether HSC71 acetylation-mediated ROS production represented a conserved defense mechanism, inhibitor application was conducted on *Drosophila* S2 cells. IP analysis revealed that endogenous HSPA8 acetylation increased upon EX-527 treatment, but not SirReal2 treatment (Fig 6G), suggesting that HSPA8 deacetylation was primarily mediated by SIRT1 in *Drosophila* S2 cells. EX-527 treatment also significantly increased intracellular ROS levels and SOD activity in *Drosophila* S2 cells (Fig 6H, 6I). Given our previous finding that EsSOD knockdown elevated ROS level in crab hemocytes, we overexpressed EsSOD and DmSOD in *Drosophila* S2 cells to validate the role of SOD in ROS regulation. As expected, both EsSOD and DmSOD overexpression reduced intracellular ROS levels (Fig 6J). EX-527 treatment partially restored intracellular ROS levels that were suppressed by DmSOD overexpression (Fig 6K). IP analysis further revealed that EX-527 impaired the interaction between endogenous HSPA8 and DmSOD, likely by disrupting their binding through HSPA8 hyperacetylation (Fig 6L, 6M). Moreover, following *S. eriocheiris* infection, DmSOD overexpression significantly reduced the viability of *Drosophila* S2 cells and increased the intracellular *S. eriocheiris* load. Administration of EX-527 partially rescued *S. eriocheiris* infection-induced cell death and suppressed pathogenic proliferation (Fig 6N, 6O). Taken together, EX-527 exhibited a conserved protective effect against *S. eriocheiris* in both crabs and the invertebrate model *Drosophila* S2 cells by promoting HSC71 acetylation and ROS production.

## Discussion

The HSP70 protein family plays critical roles in host-pathogen interactions, primarily by modulating viral entry, intracellular trafficking, and replication, as well as activating host innate immune responses [32, 33]. For instance, HSP70 interacted with the PB2 and PB1 subunits of the influenza A virus polymerase complex, shuttling them into the nucleus to enhance viral transcription and replication [34]. Similarly, during KSHV infection, BiP (HSPA5), an ER-resident Hsp70 family member, was upregulated and supported viral lytic replication by promoting the survival of infected cells and ensuring efficient production of viral particles [35]. HSP70 could also bind to bacterial lipopolysaccharide (LPS), facilitating signal delivery to CD14 and Toll-like receptor clusters, thereby amplifying inflammatory responses [36]. Additionally, HSPA8 prevented K48-mediated ubiquitination and degradation of RHOB and BECN1, promoted the assembly of the HSPA8-RHOB-BECN1 complex via liquid-liquid phase separation, and thus facilitated autophagosome formation and the clearance of intracellular bacteria [37]. In this study, HSC71 was significantly induced and concurrently underwent deacetylation upon *S. eriocheiris* infection. Knockdown of HSC71 reduced crab survival and promoted *S. eriocheiris* propagation in hemocytes, providing further evidence for the involvement of HSC71 in resistance to microbial infection. Additionally, it is well established that HSP70 acts as a negative regulator in apoptosis. By binding to pro-apoptotic factors such as AIF and Apaf-1, HSP70 blocks Apaf-1-mediated apoptosome formation and sequesters AIF to inhibit nuclear chromatin condensation, which ultimately suppresses the apoptotic cascade [38, 39]. Likewise, RNAi-mediated HSP70 knockdown has been shown to downregulate the anti-apoptotic protein Bcl-X_L_ and promotes caspase-3-dependent apoptosis in erythroid precursors [40]. Inhibition of the mitochondrial HSP70 Mortalin with the small-molecule inhibitor Az-TPP-O3 disrupted its interaction with p53, resulting in mitochondrial outer membrane permeabilization and the induction of mitochondrial-dependent apoptosis [41]. Consistent with these findings, HSC71 deficiency in our study triggered apoptosis in hemocytes, demonstrating a conserved anti-apoptotic function for HSC71 in crabs.

Acetylation is a common post-translational modification that regulates protein function, such as enzymatic activity, stability, subcellular localization and interactions [42]. As a ubiquitous molecular chaperone present in diverse cellular compartments, HSP70 is involved in various cellular processes through its interaction with cofactors [5]. Previous study has shown that acetylation of HSP70 at K126 enhanced its sequestration of Bcl2, thereby promoting autophagosome formation and mitophagy in MDA-MB-231 cells [43]. Conversely, a K77R mutation, which mimics an acetylation-deficient state, inhibited the interaction of HSP70 with AIF and Apaf-1, leading to caspase-3 activation and apoptosis [44]. Our previous study demonstrated that infection with *S. eriocheiris* induced apoptosis and cytopathic effect in *Drosophila* S2 cells [30]. In the present study, overexpression of hyperacetylation-mimetic HSC71(K579Q) conferred stronger protection to *Drosophila* S2 cells against *S. eriocheiris* induced-apoptosis in *Drosophila* S2 cells than wild-type HSC71, suggesting that acetylation of HSC71 represented a regulatory mechanism to enhance host cell survival.

Acetylation is catalyzed by lysine transferases (KATs), which transfer an acetyl group from acetyl-CoA to the ε-amino group of lysine residues; this process is reversed by lysine deacetylases (KDACs). It is estimated that over 90% of known acetylated proteins are primarily modified by five canonical KATs: p300, CBP, PCAF, GCN5 and Tip60 [42]. Beyond these, a growing number of non-canonical KATs have been reported to mediate protein acetylation, including ACAT1 and ARD1 [20, 45]. Carnitine acetyltransferase (Crat) is a mitochondrial enzyme to regulate the reversible transfer of acyl groups between acyl-CoA and carnitine, which has been previously linked to metabolic processing and protein acetylation [46, 47]. In human ovarian cancer cells, Crat knockdown was shown to reduce the acetylation of PGC-1α and enhance mitochondrial biogenesis, thereby promoting mitochondrial metabolism [48]. Moreover, acetylproteomic analyses have revealed that Crat deficiency also led to reduced acetylation levels of non-mitochondrial proteins [49]. In this study, Crat was identified as a acetyltransferase that modified HSC71 at K579. This modification subsequently inhibited the ubiquitination and degradation of HSC71. Consequently, Crat knockdown increased apoptosis in hemocytes and rendered crabs more susceptible to *S. eriocheiris* infection, indicating that Crat played an important role in regulating innate immunity. Moreover, the K579 residue was strictly conserved across diverse crustaceans but absent in representative vertebrates and insects. This phylogenetic conservation suggested that it might constitute a critical point of functional differentiation, with its acetylation likely representing a key functional innovation.

The crosstalk between acetylation and ubiquitination is increasingly recognized as a widespread regulatory mechanism, frequently orchestrated by specific acetyltransferases to dynamically control protein stability. MOB1 was acetylated at K11 by acetyltransferase CBP, and stabilized itself through limiting the binding affinity for E3 ligase Praja2 and subsequent ubiquitination [50]. Similarly, acetyltransferase GCN5 could acetylate DACH1 at K640, which enhanced its interaction with deubiquitinase USP7, ultimately resulting in DACH1 deubiquitination and stabilization [51]. Furthermore, our results demonstrated that overexpression of Crat in *Drosophila* S2 cells increased the acetylation and expression levels of HSC71 and simultaneously inhibiting its ubiquitination. These observations led us to hypothesize that Crat might interfere with the binding of an E3 ubiquitin ligase to HSC71 and thus regulated protein stability. To test this hypothesis, we co-expressed HSC71 and Crat with CHIP, a well-characterized E3 ubiquitin ligase for the HSC70 family [31]. Strikingly, Crat overexpression attenuated the strong interaction between CHIP and HSC71, thereby suppressing the CHIP-induced increase in HSC71 ubiquitination and restoring it to near-basal levels, which consequently prevented CHIP-mediated degradation of HSC71. These results revealed a novel mechanism whereby Crat regulated HSC71 stability by blocking CHIP binding.

CHIP is a co-chaperone of HSP70/HSC70, containing a C-terminal U-box domain with ubiquitin ligase activity that targets proteins for degradation. Through its N-terminal tetratricopeptide repeat (TPR) domain, CHIP binds to the C-terminal lid subdomain and the conserved EEVD motif of HSC70 [52, 53]. This binding enables CHIP to facilitate the ubiquitination and proteasomal degradation of HSC70 and client proteins chaperoned by HSC70 [31, 54]. Thus, the regulation of the CHIP-HSC70 interaction is a key determinant in intracellular protein quality control, governing the switch between protein refolding and degradation. Notably, acetylation of HSP70 at K77 by ARD1 was crucial for maintaining its chaperone function. The K77R mutation in the NBD of HSP70 reduced its affinity for the co-chaperones HOP and HSP90. This reduced binding preference shifted HSP70 towards the E3 ubiquitin ligase CHIP, leading to enhanced HSP70 ubiquitination and degradation, and ultimately promoting apoptosis under cellular stress [20]. In this study, the acetylation site K579 of HSC71 was located within its substrate-binding domain (SBD), a region known to directly interact with CHIP. It was therefore plausible that the K579R mutation weakened the CHIP-HSC71 interaction, potentially by inducing a conformational change, which consequently reduced HSC71 ubiquitination. In addition to allosteric modulation of chaperone activity, the CHIP-HSP70 interaction was also regulated through competitive binding with another E3 ubiquitin ligase, Abs10. Unlike CHIP, Abs10 did not directly catalyze HSP70 ubiquitination but rather antagonized CHIP-mediated ubiquitination and degradation of HSP70 [55]. In the present study, a distinct interaction between Crat and HSC71 was observed in *Drosophila* S2 cells upon CHIP co-expression. This observation prompted us to hypothesize that Crat binding likely occupied the CHIP-binding site on HSC71, thereby suppressing CHIP’s ubiquitin ligase activity toward HSC71 and promoting HSC71 stabilization.

ROS induce oxidative damage that is detrimental to intracellular microbes, thereby serving as an effective antimicrobial defense for the host [56]. The most physiologically relevant ROS include the superoxide anion (O_2_ ^•−^) and hydrogen peroxide (H O), which are generated sequentially. O_2_ ^•−^ is primarily produced by specific enzymes such as NOXs and is detoxified by SODs into oxygen (O_2_) and H_2_O_2_. Subsequently, H_2_O_2_ is decomposed by catalases, glutathione peroxidases, and peroxiredoxin, collectively maintaining cellular ROS homeostasis [57]. In our study, several antioxidant proteins, including peroxiredoxin and SOD, were identified as HSC71 interactors. Given the established roles of peroxiredoxin and SOD in scavenging ROS and their interactions with HSP70, it is plausible that HSC71 could indirectly influence ROS homeostasis [58–60]. Notably, overexpression of the K579 mutant significantly restricted *S. eriocheiris* propagation in *Drosophila* S2 cells, an effect accompanied by elevated intracellular ROS levels. Consistent with a role for acetylation in this process, knockdown of the acetyltransferase Crat reduced both HSC71 acetylation and intracellular ROS levels. Both the K579Q mutation and Crat overexpression attenuated the binding of HSC71 to SOD, whereas the K579R mutation enhanced this interaction. In contrast, neither the K579Q nor the K579R mutation affected the association between HSC71 and peroxiredoxin. These indicated that the acetylation-dependent regulation of the HSC71-SOD interaction specifically modulates ROS production, thereby contributing to the limitation of *S. eriocheiris* infection.

SIRT1, an NAD^+^-dependent histone deacetylase, participates in diverse cellular processes such as stress response, metabolism and tumorigenesis [61]. It targets a broad spectrum of acetylated proteins, with over 40% of acetylation sites serving as SIRT1 substrates [42]. Notably, SIRT1 exerts sophisticated control over the HSP70 chaperone network. For example, SIRT1 deacetylated heat shock 70 kDa protein 4 (HSPA4) at lysine residues K305, K351 and K605, promoting its nuclear translocation and thereby suppressing proinflammatory cytokine expression [62]. Similarly, deacetylation of heat shock factor 1 (HSF1) at K80 by SIRT1 prolonged HSF1 binding to the heat shock promoter of HSP70, leading to enhanced HSP70 expression [63]. Moreover, SIRT1 also contributed to the maintenance of redox homeostasis by deacetylating the transcription factor Foxo3. This modification upregulated the expression of antioxidant enzymes, including CAT and manganese superoxide dismutase (MnSOD), and thus alleviated mitochondrial oxidative stress [64]. Consistently, icariin could activate SIRT1, which upregulated the transcriptional cofactor PGC-1α, thereby restoring the activity of endogenous antioxidant enzymes like SOD and glutathione peroxidase to mitigate subarachnoid hemorrhage-induced oxidative stress. Importantly, this antioxidant effect was abolished by the selective SIRT1 inhibitor EX-527, confirming the specificity of the SIRT1-dependent mechanism [65]. In our study, the acetylation levels of both crab HSC71 and its *Drosophila* homolog HSPA8 were elevated upon treatment with EX-527, indicating that SIRT1 was a deacetylase responsible for HSC71 deacetylation. Furthermore, EX-527 administration significantly increased ROS production in crab hemocytes and *Drosophila* S2 cells, consistent with our earlier finding that the hyperacetylation-mimetic HSC71(K579Q) elevated ROS levels. Importantly, in the invertebrate model *Drosophila* S2 cells, EX-527 treatment attenuated the interaction between endogenous HSPA8 and DmSOD. This disruption is attributable to the compromised chaperone activity of HSPA8 resulting from EX-527-induced hyperacetylation, which consequently impaired its ability to form a stable complex with DmSOD. While HSP70 was known to prevent SOD aggregation and facilitate SOD’s mitochondrial transport for ROS scavenging, it could also target SOD for CHIP-mediated degradation [58, 66]. Our data, however, showed that modulating HSC71 acetylation did not affect SOD stability, indicating that the HSC71-SOD interaction was uncoupled from proteasomal turnover. Instead, we proposed a model in which acetylation at HSC71 K579 acted as a switch: in its deacetylated state, HSC71 assisted in delivering SOD to mitochondria to clear ROS; acetylation abrogated this chaperone function, leading to ROS accumulation, which in turn restricted intracellular *S. eriocheiris* proliferation.

In conclusion, this study established a critical role for HSC71 acetylation in regulating host defenses against *S. eriocheiris* infection (Fig 7). Specifically, the acetylation level of HSC71 was consistently reduced following infection with *S. eriocheiris*. HSC71 knockdown aggravated hemocyte apoptosis and *S. eriocheiris* invasion in crabs, whereas hyperacetylation-mimetic HSC71(K579Q) overexpression suppressed *S. eriocheiris* infection-induced apoptosis and improved survival in *Drosophila* S2 cells. Mechanistically, Crat functioned as the acetyltransferase responsible for acetylating HSC71 at K579, and counteracting CHIP-mediated ubiquitination and degradation of HSC71. Loss of Crat reduced the acetylation and expression levels of HSC71, thereby increasing hemocyte apoptosis and crab susceptibility to *S. eriocheiris* infection. Furthermore, acetylation at K579 disrupted the interaction between HSC71 and SOD, leading to diminished SOD activity and elevated ROS production. Treatment of crabs and *Drosophila* S2 cells with the SIRT1 inhibitor EX-527 elevated endogenous HSC71 acetylation and increased intracellular ROS levels, thereby enhancing resistance to *S. eriocheiris* infection. Therefore, we uncovered a previously unrecognized mechanism by which Crat-mediated acetylation of HSC71 stabilized the protein and modulated ROS homeostasis to orchestrate an effective innate immune defense against *S. eriocheiris* infection. Our findings provided a potential strategy of targeting HSC71 acetylation to intervene in *S. eriocheiris* infection.

**Fig. 7.**
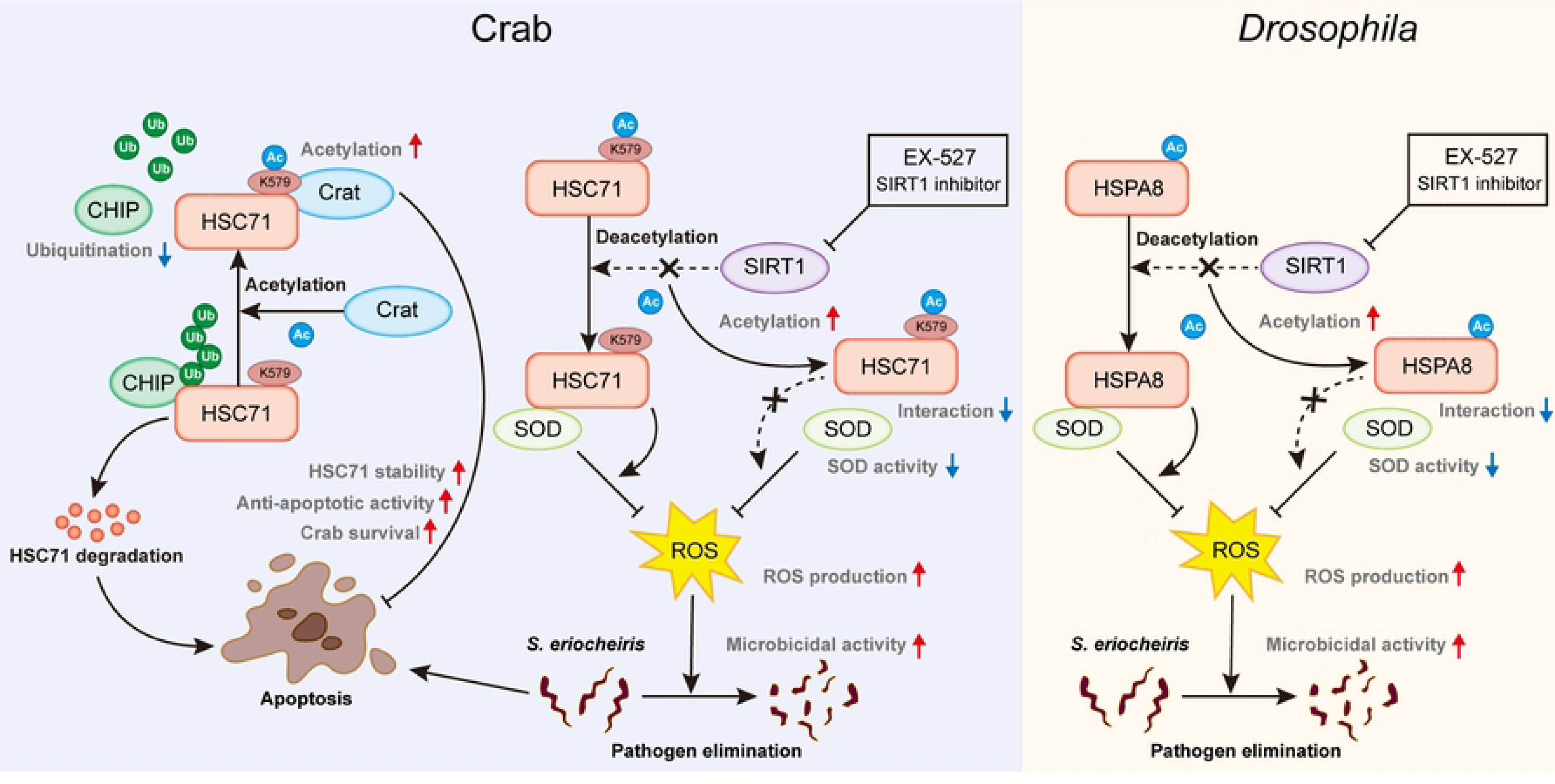
A proposed model for HSC71 acetylation-mediated resistance to *S. eriocheiris* infection in arthropods. HSC71 is an anti-apoptotic protein that protects crabs against *S. eriocheiris* infection. Crat-mediated HSC71 acetylation at K579 weakens the interaction between HSC71 and CHIP, inhibiting HSC71 ubiquitination and degradation, which enhances anti-apoptotic capacity to defend *S. eriocheiris* infection. Concurrently, K579 acetylation impairs the binding of HSC71 to SOD, leading to decreased SOD activity, accompanied by increased ROS levels. As a selective SIRT1 inhibitor, EX-527 elevates HSC71 acetylation and enhances ROS production in hemocytes, subsequently inhibiting *S. eriocheiris* infection of crabs. Additionally, EX-527 also increases HSPA8 acetylation and ROS levels in *Drosophila* S2 cells, and thus restricts the proliferation of *S. eriocheiris*, indicating a conserved protective mechanism across arthropods.

## Materials and methods

### Experimental animal and pathogen

Chinese mitten crab (*Eriocheir sinensis*, approximately 25 g) were obtained from an aquaculture farm in Baoying, Jiangsu Province, China, and acclimated for one week at 28℃ in a laboratory recirculating aquaculture system before experiments. *S. eriocheiris* isolated from crabs with “tremor disease” [67] was cultured in R2 medium at 30℃ for 16 h. The bacterial pellets were harvested by centrifugation at 12,000×g for 10 min and resuspended in sterile PBS. For *in vivo* infection, each crab was injected with 50 μL of *S. eriocheiris* suspension (4×10^8^ cells/mL). For *in vitro* infection, *Drosophila* S2 cells seeded in 24-well or 96-well plates were infected with *S. eriocheiris* at a multiplicity of infection (MOI) of 30 for 48 h [30].

### Plasmids, antibodies and reagents

Full-length HSC71 was amplified from the cDNA of *E. sinensis* hemocytes and cloned into pAc5.1-GFP and pAc5.1-HA vectors. Similarly, full-length carnitine O-acetyltransferase (Crat), superoxide dismutase (SOD) and peroxiredoxin (Prx) were cloned into pAc5.1 vectors to generate GFP-, RFP- or FLAG-tagged Crat, SOD and Prx plasmids. Full-length CHIP was cloned into pAc5.1-V5 and pAc5.1-GFP vectors. HA-Ub was generated by cloning ubiquitin into pAc5.1-HA vectors. The *Drosophila* SOD homolog (DmSOD) was amplified from the cDNA of *Drosophila* S2 cells and cloned into pAc5.1-FLAG vectors. Mutant plasmids expressing HSC71-GFP (K579Q) and HSC71-GFP (K579R) were generated by mutagenesis using the Mut Express II Fast Mutagenesis Kit (C214, Vazyme). All constructs were confirmed by DNA sequencing. All primer sequences used for cloning and mutagenesis were provided in S3 Table.

Antibodies used in this study were listed as follows, anti-Acetyllysine (105RM, PTM, China), anti-HSPA8 (MH4829, Abmart, China), anti-β-actin (HC201, Transgen, China), anti-GFP (HT801, Transgen, China), anti-Tubulin (HC101, Transgen, China), anti-HA (HT301, Transgen), anti-Tubulin (HC101, Transgen, China), anti-FLAG (HT201, Transgen, China), anti-V5 (AF2894, Beyotime, China), anti-Ubiquitin (AG3161, Beyotime, China), and mouse IgG (A7028, Beyotime, China). Anti-GFP (P2132) and anti-FLAG (P2115) magnetic beads were purchased from Beyotime. Protein A/G magnetic beads (PB101) were purchased from Vazyme. The proteasome inhibitor MG132 (S1748), SIRT1 inhibitor EX-527 (SC0281; 10 mM in DMSO) and SIRT2 inhibitor SirReal2 (SC0290; 10 mM in DMSO) were purchased from Beyotime. The SIRT family deacetylase inhibitor Nicotinamide (NAM, 51402ES03) and HDAC family deacetylase inhibitor Trichostatin A (TSA, 51406ES03) were purchased from Yeason.

### Cell culture and transfection

*Drosophila* S2 cells (R69007, Gibco, USA) were cultured at 28℃ in serum-free insect cell medium (12658027, Gibco, USA). Primary hemocytes were isolated from crab hemolymph as previously reported [68] and maintained at 28℃ in Leibovitz’s L-15 medium (11415064, Gibco, USA). Plasmid transfection in *Drosophila* S2 cells was performed using FuGENE HD transfection reagent (E2311, Promega, USA) according to the manufacturer’s instructions.

## Acetylome

Hemocytes from *S. eriocheiris*-infected and healthy crabs were lysed in urea buffer (8 M Urea, 100 mM Tris-HCl, pH 8.5) and digested using filter-assisted sample preparation (FASP). The resulting peptides were lyophilized and subjected to acetylated peptide enrichment with the PTMScan Acetyl-Lysine Motif [Ac-K] Kit (Cell Signaling Technology, 13416) according to the manufacturer’s instructions. Peptides were separated on a homemade C18 column using a 60-min linear gradient (buffer A: 0.1% FA; buffer B: 84% ACN, 0.1% FA) at 300 nL/min on a NanoElute system coupled to a timsTOF Pro mass spectrometer (Bruker Daltonics, USA). MS data were acquired in positive ion mode over m/z 100-1700 and 1/k0 0.6-1.6, with PASEF MS/MS (10 cycles; target: 1.5k; threshold: 2.5k) and 0.4-min active exclusion. The raw data were processed in MaxQuant for identification and quantification. Hierarchical clustering was performed using Cluster 3.0 and visualized with Java TreeView software. Enrichment analysis was conducted using Fisher’s exact test (with the full quantified proteome as background), and a Benjamini-Hochberg adjusted *p*-value < 0.05 was considered statistically significant.

## Quantitative real-time PCR and absolute quantitative PCR

Total RNA extraction was performed using FastPure Tissue/Cell Total RNA isolation Kit (RC113, Vazyme, China). Reverse transcription of the extracted RNA into cDNA utilized HiScript III 1^st^ Strand cDNA Synthesis Kit (R312, Vazyme, China). Quantitative real-time PCR was performed using ChamQ SYBR qPCR Master Mix (Q311, Vazyme, China) on a LightCycler 96 System (Roche, Germany). GAPDH was used for normalization, and the relative expression of target genes was calculated using the 2^−ΔΔCT^ method.

Genomic DNA was extracted from infected crab hemocytes or *Drosophila* S2 cells using EasyPure Genomic DNA Kit (EE101, Transgen, China) for quantification of *S. eriocheiris* load. Absolute quantitative PCR was performed as detailed in our previous study [69]. The *S. eriocheiris* copy number was determined using a standard curve generated from serial dilutions of a plasmid containing a *S. eriocheiris* genomic fragment. All primer sequences used in this study were shown in S3 Table.

## Cell viability assay

*Drosophila* S2 cells were seeded in triplicate into 96-well plates and incubated for 12 h before transfection or inhibitor treatment. Following challenge with *S. eriocheiris* for 48 h, cell viability was assessed by adding CCK-8 (A311, Vazyme, China) to a final concentration of 10% and incubating for 1 h. Relative cell viability was calculated by normalizing the absorbance at 450 nm of wells containing cells treated as indicated to that of untreated control wells.

## RNA interference

To knock down the expression of HSC71 and Crat, gene-specific siRNAs were commercially synthesized (S3 Table). Each crab was injected with 20 μg of the respective siRNA, with crabs received an equivalent volume of non-targeting control siRNA as controls. Knockdown of SOD was performed by application of double-stranded RNA (dsRNA) *in vivo*. Primers with T7 promoters (S3 Table) were used to amplify the partial DNA fragments of SOD. The T7 RNA polymerase (EP0111, Thermo, USA) was employed to synthesize dsRNA specific for SOD according to the manufacturer’s instructions, with dsRNA targeting GFP produced similarly as controls. The synthesized dsRNA was dissolved in DEPC water to a final concentration of 1 μg/μL. Each crab received 40 μL of the solution, followed by a second injection of the same dose 24 h later. Hemocytes were collected at 24, 48 and 72 h after siRNA injection, or 48, 72 and 96 h after the initial dsRNA injection. Knockdown efficiency of each gene was assessed by quantitative real-time PCR. Following validation of the RNA interference effect, subsequent experiments were conducted 24 h after siRNA injection or 48 h after the initial dsRNA injection.

## Mass spectrometry for identification of HSC71 interactors

Hemocytes from healthy crabs were lysed on ice for 30 min in lysis buffer (20 mM Tris-HCl pH 7.5, 150 mM NaCl, 1% Triton X-100 and protease inhibitors). Cell lysates were centrifuged at 12,000×g for 10 min, and the supernatants were incubated overnight at 4℃ with protein A/G magnetic beads conjugated with an anti-HSPA8 antibody for immunoprecipitation (IP). Beads coated with non-specific mouse IgG were used as a negative control. After washing three times with lysis buffer, precipitated proteins were eluted from the beads with SDT buffer (4% SDS, 100 mM Tris-HCl, 1 mM DTT, pH 7.6). The elution was visualized by Coomassie staining and subsequently analyzed by liquid chromatography-tandem mass spectrometry (LC-MS/MS) on a Q Extractive mass spectrometer (Thermo, USA).

## Co-immunoprecipitation and immunoblotting

*Drosophila* S2 cells were transfected with the indicated plasmids for 48 h. After specific treatment, cells were lysed in IP lysis buffer (P0013, Beyotime, China) containing protease inhibitor cocktail (P1005, Beyotime, China) and deacetylase inhibitor cocktail (P1112, Beyotime, China). Cell lysates were subjected to IP using magnetic beads conjugated with specific antibodies for 12 h at 4℃. The precipitates were washed three times with IP lysis buffer, and the immune complex were eluted by boiling in protein buffer for immunoblotting analysis.

For immunoblotting, protein samples were separated by SDS-PAGE and transferred onto polyvinylidene fluoride membranes. After blocking with 5% silk milk for 2 h at room temperature, the membranes were incubated with primary antibodies as mention above overnight at 4℃, followed by incubation with corresponding secondary antibodies for 1 h at room temperature. The immunoreactive bands were visualized using a chemiluminescence imaging system (Tanon, China) and quantified with Image J software. Uncropped western blots were shown in S3-S11 Fig.

## Apoptosis assay

Hemocytes after RNA interference were harvested by centrifugation at 600×g for 5 min and washed twice with PBS. Apoptosis in hemocytes was assessed by staining with Annexin V-FITC and PI using the Apoptosis Detection Kit (C1062, Beyotime, China) following the manufacturer’s instructions. *Drosophila* S2 cells overexpressing wild-type HSC71-GFP or mutants were infected with *S. eriocheiris* for 48 h. Apoptosis was then assessed using Annexin V-PE (C1065, Beyotime, China) instead of Annexin V-FITC to avoid spectral overlap with GFP fluorescence. Untransfected cells were used to define the gating strategy for analyzing transfected populations. The apoptotic cells were analyzed on a FACSVerse flow cytometer (BD, USA), and the apoptotic rates were quantified using FlowJo software as indicated in the respective figure legends. For mitochondrial membrane potential detection, hemocytes following RNA interference were processed using the Mitochondrial Membrane Potential Assay Kit with Rhodamine 123 (C2008S, Beyotime, China) according to the manufacturer’s instructions. Ten thousand cells per sample were analyzed on the flow cytometer, and the mean fluorescence intensity was quantified using FlowJo software.

## Assessment of intracellular ROS

For intracellular ROS detection, hemocytes and *Drosophila* S2 cells were rinsed three times with sterile PBS and stained with 10 μM DCFH-DA (S0033S, Beyotime, China) at 28℃ for 30 min. Meanwhile, *Drosophila* S2 cells overexpressing wild-type HSC71-GFP or mutants were stained with 5 μM dihydroethidium (DHE, 50102ES02, Yeason) at room temperature for 1 h. After the ROS probe staining, the cells were washed three times with sterile PBS and analyzed using a FACSVerse flow cytometer (BD, USA). For each analysis, ten thousand events were acquired, and the mean fluorescence intensity was quantified using FlowJo software.

## Measurement of SOD activity

Following transfection or inhibitor treatment, hemocytes or *Drosophila* S2 cells were harvested and lysed on ice by ultrasonication. Protein concentration was determined using a Bradford protein assay kit (P0006, Beyotime, China), while SOD activity was quantified with a superoxide dismutase assay kit (A001-3-2, Jiancheng, China) according to the manufacturer’s instructions.

## Inhibitor application

For *in vitro* experiments, hemocytes were treated with NAM (5 mM) or TSA (3 μM) 12 h before IP to assess HSC71 acetylation. *Drosophila* S2 cells were pretreated with EX-527 or SirReal2 at a final concentration of 1 μM for 24 h. The cells were then analyzed for HSC71 acetylation, ROS levels and antioxidant capacity, and were subsequently challenged with *S. eriocheiris*. For *in vivo* experiments, crabs were injected with 20 μL of a solution containing EX-527 or SirReal2, administered as 5 or 10 μL aliquots of a 10 mM stock diluted in DMSO. Hemocytes were isolated 6 h later to evaluate HSC71 acetylation, ROS levels and antioxidant capacity. For survival analysis, crabs received EX-527 (0.5 μg/g) 6 h before *S. eriocheiris* inoculum, with the same dose of EX-527 administered at 4 and 8 d post-infection to sustain inhibition.

## Statistical analysis

Data from at least three biological replicates were presented as the mean ± standard deviation (SD). Statistical significance for comparisons between two groups was assessed using a two-tailed Student’s t-test. Survival curves were generated by the Kaplan-Meier method, and two-group datasets were compared by the log-rank test. All statistical analyses were performed using GraphPad Prism software. Significant differences were shown by \**P*<0.05, \*\**P*<0.01, \*\*\**P*<0.001.

## Data availability statement

The acetyl-proteomics data generated in this study have been deposited in the ProteomeXchange Consortium via the iProX repository with the dataset identifier PXD 069422 [https://www.iprox.cn/page/project.html?id=IPX0013736000]. Source data are available within this article and its supplementary information files, or from the corresponding authors on request.

## Acknowledgements

Fundings

The current work was supported by grants from the “JBGS” Project of Seed Industry Revitalization in Jiangsu Province of China (No. JBGS [2021]031), and the Modern Agricultural Industry Technology System Project of Jiangsu Province of China (Grant No. JATS [2023]311).

## Author contributions

**Yubo Ma:** Conceptualization, Data curation, Investigation, Methodology, Visualization, Writing-original draft. **Xiang Meng, Xin Yin, and Yu Yao:** Investigation, Methodology. **Siyu Lu:** Formal analysis, Visualization. **Wei Gu:** Validation. **Qingguo Meng:** Funding acquisition, Supervision, Writing-review & editing.

## Supporting information

S1 Table. Identification and quantification data for acetylated peptides from LC-MS/MS.

S2 Table. Raw data from LC-MS/MS analysis of immunoprecipitated proteins. S3 Table. Primer and siRNA sequences used in this study.

S1 Fig. Bioinformatics analysis of differentially acetylated proteins in response to *S. eriocheiris* infection. (A and B) Gene ontology (GO) annotation and enrichment analysis of differentially acetylated proteins. (C-E) Kyoto Encyclopedia of Genes and Genomes (KEGG) pathway annotation and enrich analysis of differentially acetylated proteins.

S2 Fig. Sequence information of EsHSC71. (A) Nucleotides and amino acids of EsHSC71. (B) Alignment of multiple amino acid sequences between EsHSC71 and its homologs from other species.

S3 Fig. Uncropped western blots for Fig 1C, Fig 1E, Fig 2D, Fig 2E. S4 Fig. Uncropped western blots for Fig 3B, Fig 3C, Fig 3D, Fig 3E.

S5 Fig. Uncropped western blots for Fig 4C, Fig 4E. S6 Fig. Uncropped western blots for Fig 4F, Fig 4G. S7 Fig. Uncropped western blots for Fig 4H, Fig 4I.

S8 Fig. Uncropped western blots for Fig 4J, Fig 4K, Fig 4L, Fig 4M. S9 Fig. Uncropped western blots for Fig 5B, Fig 5C, Fig 5D.

S10 Fig. Uncropped western blots for Fig 5H, Fig 6A, Fig 6B, Fig 6G. S11 Fig. Uncropped western blots for Fig 6L, Fig 6M.

